# Differential substrate affinity and catabolite repression enable preferential use of urea by ammonia-oxidizing bacteria

**DOI:** 10.1101/2023.08.04.551995

**Authors:** Wei Qin, Stephany P. Wei, Yue Zheng, Eunkyung Choi, Xiangpeng Li, Juliet Johnston, Xianhui Wan, Britt Abrahamson, Zachary Flinkstrom, Baozhan Wang, Hanyan Li, Lei Hou, Qing Tao, Wyatt W. Chlouber, Xin Sun, Michael Wells, Long Ngo, Kristopher A. Hunt, Hidetoshi Urakawa, Xuanyu Tao, Dongyu Wang, Xiaoyuan Yan, Dazhi Wang, Chongle Pan, Peter K. Weber, Jiandong Jiang, Jizhong Zhou, Yao Zhang, David A. Stahl, Bess B. Ward, Xavier Mayali, Willm Martens-Habbena, Mari-Karoliina H. Winkler

## Abstract

Four distinct lineages of ammonia-oxidizing microorganisms (AOM) collectively contribute to one of the largest nitrogen fluxes in the global nitrogen budget. AOM possess widely different specific affinities for ammonia, thought to determine their niche differentiation. Nevertheless, ammonia-oxidizing archaea and bacteria (AOA, AOB), and complete ammonia oxidizers (comammox) co-occur in soils, freshwater sediments, and aquifers, suggesting that other factors must drive their coexistence. Here, we show that representatives of four AOM lineages employ distinct regulatory strategies for ammonia or urea utilization, thereby minimizing direct competition for either substrate. The tested AOA and comammox species preferentially used ammonia over urea, while beta-proteobacterial AOB favored urea utilization, repressed ammonia transport in the presence of urea, and showed higher affinity for urea than ammonia, whereas gamma-proteobacterial AOB co-utilized both substrates. Stable isotope tracing, kinetics, and transcriptomics experiments revealed that both assimilation and oxidation of ammonia are transport-dependent. These results reveal novel mechanisms of nitrogen metabolism regulation and transporter-based affinity underlying the contrasting niche adaptation and coexistence patterns among the major AOM lineages.

## Introduction

Ammonia-oxidizing microorganisms (AOM) account for the oxidation of ∼2,330 Tg N per year, representing one of the largest nitrogen fluxes in the global nitrogen budget ^1^. Ammonia-**o**xidizing **b**acteria and **a**rchaea (AOB, AOA) oxidize ammonia (hereafter considered as total ammonia, NH_3_ + NH_4_^+^) to nitrite ^2, 3^, which is then oxidized to nitrate by **n**itrite-**o**xidizing **b**acteria (NOB), comprising the two-step process of nitrification. Recently, **com**plete **amm**onia **ox**idizers (comammox), capable of oxidizing ammonia to nitrate, were discovered within the NOB genus *Nitrospira* ^4, 5^. Given that all AOM use ammonia as energy and nitrogen (N) sources for growth, their specific affinity for ammonia has been considered the main determinant of their competition and niche partitioning ^6–8^. However, fundamental questions remain about the biochemical and biophysical controls of their specific affinities for ammonia ^9,10^. Although different lineages of AOM are thought to compete for their primary N substrate, ammonia, they often grow concomitantly in soil, freshwater, estuarine, and subsurface ecosystems, raising the question of other adaptative strategies that allow for their co-occurrence.

Microorganisms can maximize survival and balance growth by either utilizing multiple substrates simultaneously or preferentially using those that support optimal growth with least investment in enzyme inventory. A classic example of nutrient selection is the glucose-lactose diauxie in *Escherichia coli* based on the pioneering work of Jacques Monod in 1940s ^11^. Most subsequent studies have focused on how microorganisms metabolize carbon (C) mixtures ^12^ or mixed N sources for anabolism ^13, 14^. However, very little is known about the selection of alternative N substrates for catabolic energy conversion, which is a physiological trait unique to nitrifying microorganisms ^1^.

Many AOM species can utilize urea as an energy and N source through hydrolysis to ammonia, and such metabolic flexibility offers adaptive advantage ^4, 15–20^. Urea is an important and widely available organic nitrogen compound in soil and aquatic environments which are large components of the global nitrogen cycle. It is released as nitrogenous waste from both eukaryotes and prokaryotes and subsequently hydrolyzed by ureolytic microorganisms ^21, 22^. Urea-based fertilizers are a mainstay in modern agriculture, and the intensive use of urea in fertilizers has greatly increased urea export to surrounding freshwater and coastal ecosystems. Since the urease enzyme does not require a high-energy cofactor, such as ATP ^23^, recent studies in marine systems suggested that ammonia and urea may be co-utilized for energy and N requirements ^24^. However, utilization or even co-utilization of urea would require an investment in the synthesis of the urea transporter, urease, and accessory proteins, a significant cost for autotrophic AOM using the low energy-yielding ammonia oxidation process. Thus, there is a compelling need for better mechanistic understanding of urea utilization by ammonia oxidizers having capacity to use this substrate.

Here we use a combination of stable isotope tracing, kinetics, and transcriptomics studies to resolve the distinct regulatory strategies for uptake and metabolism of ammonia and urea by AOA, beta- and gamma-proteobacterial AOB, and comammox. Developing that understanding also offered fundamental new insights into the core biochemistry and sensory systems of any organism using reduced N species for both energy generation and biosynthesis. The preference for urea and selective repression of ammonia transport by the tested β-AOB species indicates that, in addition to ammonia assimilation, ammonia oxidation and affinity are also transport-dependent. This was further confirmed by the adaptive regulation of cellular affinities of marine AOA and comammox for ammonia upon ammonia exhaustion. These new findings significantly advance understanding of niche separation and resource partitioning among lineages of AOM and should foster improved nitrogen cycle models.

## Results

### Different AOM lineages exhibit distinct N source preferences for energy metabolism

Comparative analysis of available AOM genomes showed that more than 50%, 60%, and 80% of AOA, AOB, and comammox species, respectively, contain genes encoding urea permease, urease, and accessory proteins (Extended Data Fig. 1). Thus, the potential for urea uptake and utilization is widespread among AOM. Seven phylogenetically and ecologically diverse AOM species, including a marine AOA (*Nitrosopumilus piranensis* D3C) ^15^, a soil AOA (*Nitrososphaera viennensis* EN76) ^16^, three β-AOB (*Nitrosospira lacus* APG3, *Nitrosospira multiformis* ATCC 25196, and *Nitrosomonas ureae* Nm10) ^17, 19, 20^, a marine γ-AOB (*Nitrosococcus oceani* ATCC 19707) ^18^, and a comammox (*Nitrospira inopinata* ENR4) ^4^ were selected for characterization of N substrate preference (Extended Data Fig. 2). As expected, all strains grew by near-exact stoichiometric conversion of ammonia or urea to nitrite or nitrate when grown in media containing a single N substrate (Fig. 1). No extracellular urease activity was observed for any of the strains (Extended Data Fig. 3). Additionally, neither were signal peptides identified in their urease genes, nor were secretion system genes found in the flanking regions of the urease operons, further supporting the cytoplasmic localization of urease enzymes in the tested AOM species. When grown on mixtures of equal amounts of ammonia- and urea-N, three distinct N utilization patterns emerged (Fig. 1): While the tested AOA and comammox species depleted ammonia before hydrolyzing significant amounts of urea (Fig. 1a–c), β-AOB species used urea before consuming substantial amounts of free ammonia (Fig. 1d–f). In contrast, γ-AOB species simultaneously consumed ammonia and urea (Fig. 1g).

**Figure 1.**
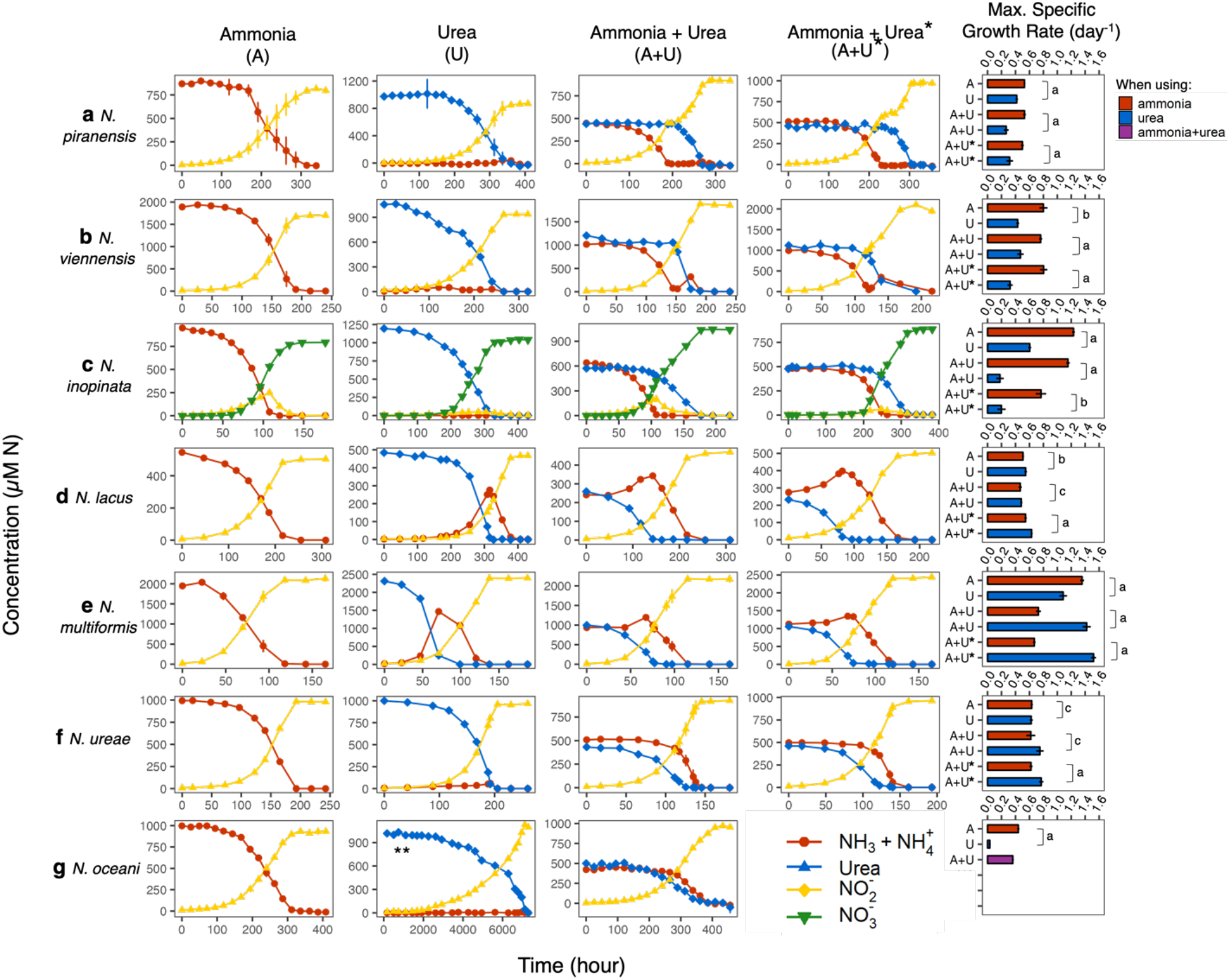
Growth curves and maximum specific growth rate (*µ_max_*) of (**a**) *N. piranensis* (marine AOA), (**b**) *N. viennensis* (soil AOA), (**c**) *N. inopinata* (comammox), (**d**) *N. lacus* (*Nitrosospira* β-AOB), (**e**) *N. multiformis* (*Nitrosospira* β-AOB), (**f**) *N. ureae* (*Nitrosomonas* β- AOB), and (**g**) *N. oceani* (γ-AOB). *Inoculation with 1% urea-grown culture at mid-exponential phase; the other group of incubations on ammonia-urea mixtures were inoculated with 1% ammonia-grown culture. Markers and error bars represent average and one standard deviation of biological triplicates. The standard deviation is smaller than the marker if error bar is not visible. **Data shown for *N. oceani*’s urea-only incubation from a single incubation without replicates. Urea-only incubations inoculated with urea-grown culture were not conducted for *N. oceani* due to the extremely slow growth rate. Letters indicate incubation with different N substrates: A = ammonia only; U = urea only; A+U = ammonia + urea. Column heights and error bars represent average and one standard deviation of biological triplicates for specific growth rate measurement. Lower case letters indicate the range of *P* values calculated from two-tailed t-test pairing. a: *P* < 0.01, b: 0.01 < *P* < 0.05, and c: *P* > 0.05. (See Supplementary Table 1 for all *P* values).

Notably, the marine AOA *N. piranensis* exhibited a marked diauxic lag between the two growth phases (Fig. 1a), reflecting a longer adaptive transition from ammonia to urea consumption. The diauxic lag phase was less pronounced for the soil AOA *N. viennensis*, displaying a rapid transition from ammonia to urea consumption (Fig. 1b). The comammox *N. inopinata* also displayed preferential consumption of ammonia when grown in the mixed N species medium (Fig. 1c). However, *N. inopinata* started to consume urea prior to the complete exhaustion of ammonia, suggesting a less stringent diauxic transition (Fig. 1c, Supplementary Discussion).

During the period of urea consumption, free ammonia concentrations increased in β-AOB *N. lacus* and *N. multiformis* (Fig. 1d and e) and decreased slightly in *N. ureae* cultures (Fig. 1f). Only after near complete consumption of urea did ammonia concentrations decrease substantially (Fig. 1d–f). The same results were observed irrespective of whether cells were pre-grown on either ammonia or urea. Unlike all other tested strains, the γ-AOB *N. oceani* co-utilized ammonia and urea without apparent preference, but only growing on urea at a rate comparable to growth on ammonia in the presence of excess ammonia in the medium (Extended Data Fig. 4). Without ammonia supplementation, *N. oceani* required almost a year to consume 500 μM urea (Fig. 1g and Extended Data Fig. 5), corresponding to an extended average generation time of 23 days.

Consistent with the preferential use of ammonia, the tested AOA and comammox species showed significantly higher specific growth rates (*P* < 0.05) on ammonia, compared to urea after ammonia exhaustion on a mixture of ammonia and urea (Fig. 1a–c). In contrast, the specific growth rates of the β-AOB strains on urea were comparable to, or even higher (*P* < 0.05) than, those on ammonia in the mixed medium (Fig. 1d–f), indicating their high urea uptake and hydrolysis activities. Together, these data indicated three contrasting N source preferences and regulation patterns among β-AOB, AOA and comammox, and γ-AOB. Whereas the tested AOA and comammox species repressed the urea-utilizing functions when ammonia was available, β-AOB activated urea transport and hydrolysis rapidly upon exposure to urea.

Although the *Nitrosospira* β-AOB strains preferentially consumed urea, release of free ammonia was observed when grown on urea or a urea-ammonia mixture (Fig. 1d and e). To evaluate the use of free ammonia in the presence of urea, and to investigate the fate of alternative N sources, we used ^15^N isotope labeling to track conversion of ammonia or urea-N throughout batch culture growth of *N. lacus*, which exhibited the most significant release of free ammonia when grown in the mixed medium (Fig. 1d). *N. lacus* was grown on a mixture of ammonia and urea, with one or the other N substrate labeled with ^15^N. For the ^15^NH_3_ and ^15^N-urea tracers, ^15^NO_2_^-^ production was measured (Fig. 2), and for the ^15^N-urea tracer, ^15^NH_3_ released to the medium was also measured (Extended Data Fig. 6, Supplementary Discussion). The temporal distribution of extracellular N isotopes exhibited three distinct phases (Fig. 2). The small amount of ^15^NO_2_^-^ produced during lag phase (0−50 h) originated mostly from ^15^NH­(Fig. 2a), reflecting metabolism of the inoculum culture grown in ammonia-only medium. However, that small amount of ^15^NO_2_^-^ was rapidly diluted by unlabeled nitrite originating from urea throughout exponential growth (∼100−150 h), reflected by the sharp drop of ^15^NH_3_ contribution to ^15^NO_2_^-^ production (Fig. 2a). Conversely, ^15^N-urea labeling showed that NO_2_^-^ almost exclusively originated from urea in this phase (Fig. 2b). In the presence of ^15^N-urea, ^15^NH_3_ was continuously released from cells until urea depletion (Extended Data Fig. 6b). During the third phase following depletion of urea around 150 h, ammonia consumption rapidly resumed, and all nitrite then originated from ammonia until the cessation of growth (Fig. 2a). Collectively, these results provided unequivocal evidence that only the ammonia released from urea hydrolysis in the cytoplasm was oxidized and that the oxidation of extracellular ammonia was repressed in the presence of urea.

**Figure 2.**
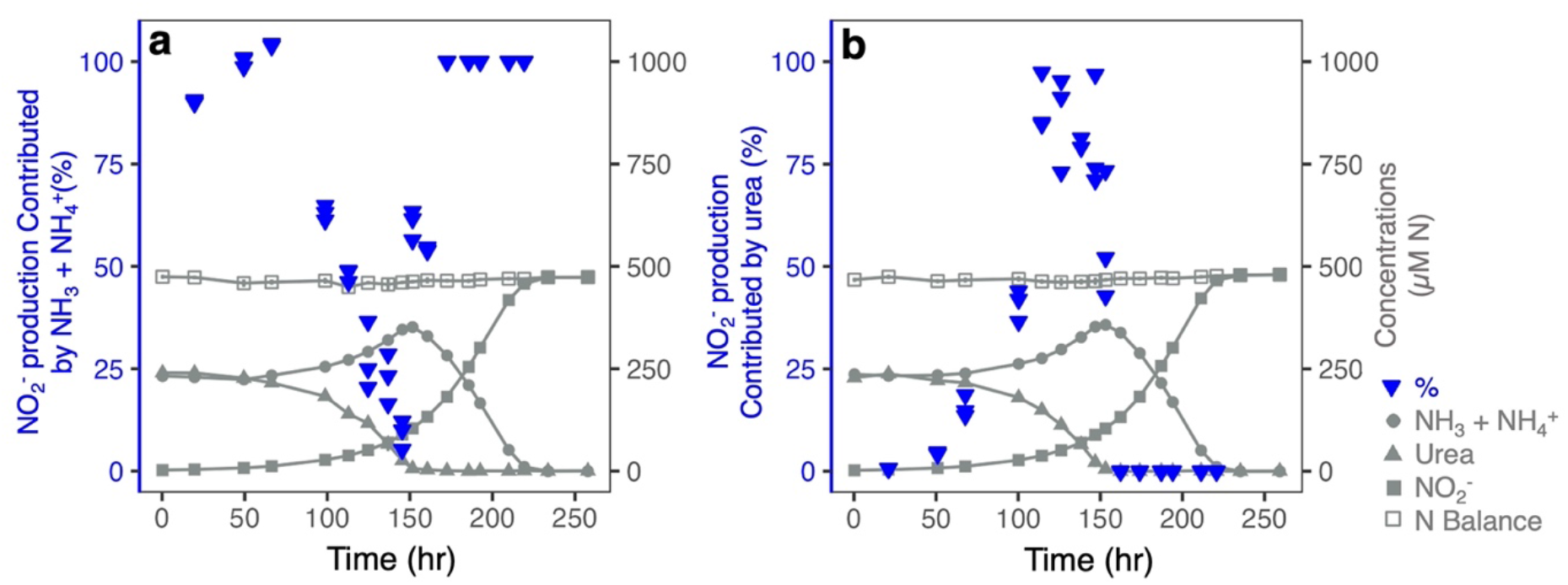
^15^N stable isotope tracking the ammonia and urea utilization in *N. lacus*. The ^15^N percentage of accumulated nitrite in the medium was measured for ^15^NH_3_ tracer incubations (**a**) and ^15^N-urea tracer incubations (**b**), in order to calculate the contribution of ammonia and urea to nitrite production. Total concentrations of N species are plotted in light grey on the secondary y axis for background reference.

### Nitrogen of preferred substrate is used for both assimilation and energy production

In addition to serving as substrates for energy conversion, ammonia- and urea-N are assimilated by AOM. We therefore examined ammonia- and urea-N assimilation at single-cell resolution in the representative AOA, β- and γ-AOB, and comammox species using stable isotope labeling in combination with NanoSIMS after growth in ammonia-urea media with either dual-labeled ^13^C^15^N-urea (plus unlabeled NH_3_ and bicarbonate) or ^15^NH_3_ and ^13^C-bicarbonate (plus unlabeled urea) (Fig. 3, Extended Data Figs. 7–10). The different AOM species cells were collected for NanoSIMS analysis during the consumption of the preferred N substrate (T1) and after the depletion of all substrates (T2) (Fig. 3a). For *N. oceani*, shown to oxidize both substrates simultaneously, cells were harvested at late exponential phase (T1) and at stationary phase (T2) after both substrates were exhausted (Fig. 3a). We also quantified assimilation of urea-derived C and compared these incorporation rates to autotrophic C fixation rates, which showed that urea is a minor source of C (Extended Data Fig. 10, Supplementary Discussion). In contrast, cells mainly assimilated urea-derived N when grown on urea.

**Figure 3.**
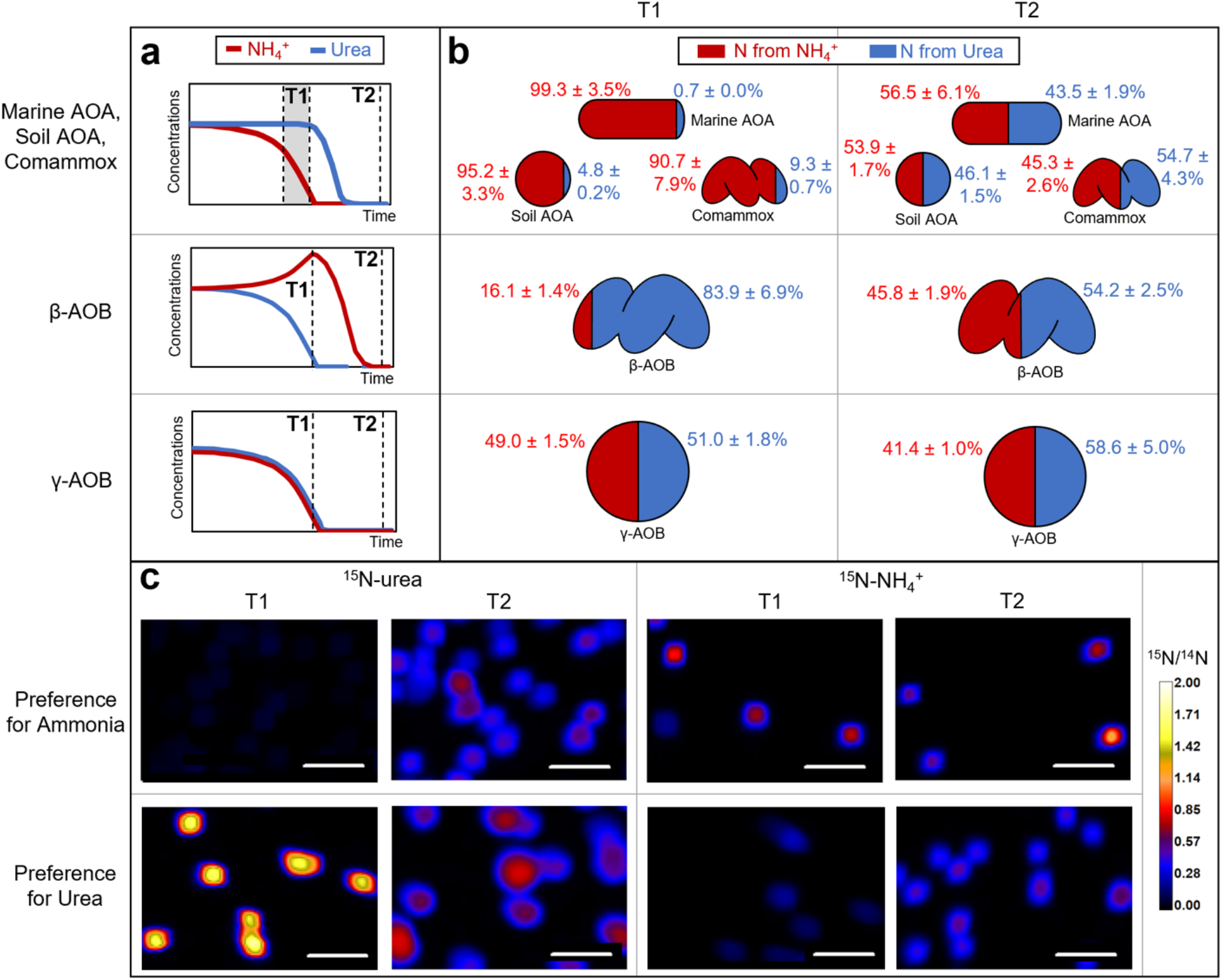
Summary of N assimilation results based on NanoSIMS. (**a**) Schematic representations of ammonia and urea consumption profiles and indication of time points for cell collection. (**b**) % new N incorporation (average ± standard deviation of all analyzed cells) from ammonia and urea. (**c**) Example NanoSIMS image (scale bar = 5 µm) where top row shows preference for ammonia (images of the soil AOA *N. viennensis* cells) and bottom shows preference for urea (images of the β-AOB *N. lacus* cells). See Extended Data Figs. 7 and 8 for growth curves with the time points indicated for each organism and the NanoSIMS images for other investigated AOM species, respectively.

Since *N. lacus* released significant ammonia into the medium when using urea, we tested whether N was assimilated from this extracellular ammonia pool or preferentially sourced from urea, even in the presence of free ammonia. Results from T1 showed 83.9 ± 6.9% new N incorporation from urea and 16.1 ± 1.4% from ammonia (Fig. 3b, c), indicating that urea-N was preferentially assimilated in the first phase of growth. Some ammonia being assimilated during lag phase is consistent with ammonia oxidation during acclimation of the ammonia-preadapted *N. lacus* to growth on a urea-ammonia mixture as discussed earlier. At T2, new N incorporation from urea had decreased to 54.2 ± 2.5% and that derived from ammonia had increased to 45.8 ± 1.9% (Figs. 3b, c). These data show that source of N incorporated by *N. lacus* is also the source of N for energy production, i.e., assimilation of urea-N was preferred, followed by ammonia assimilation after urea depletion.

A similar set of labeling experiments was performed with the marine AOA *N. piranensis*, soil AOA *N. viennensis*, and comammox *N. inopinata.* In contrast to *N. lacus,* they all preferentially assimilated N from ammonia at T1 with 99.3 ± 3.5%, 95.2 ± 3.3%, and 90.7 ± 7.9% new N incorporation from ammonia and 0.7 ± 0.0%, 4.8 ± 0.2%, and 9.3 ± 0.7% from urea for *N. piranensis*, *N. viennensis*, and *N. inopinata*, respectively (Fig. 3b, c). At T2, new N incorporation from urea and ammonia was roughly equivalent at 43.5–56.5% (Fig. 3b, c). For the third type of behavior shown by the γ-AOB *N. oceani*, we found similar new N incorporation from urea and ammonia in late exponential phase (49.0–51.0%) (Fig. 3b, c), showing that γ-AOB co-assimilated ammonia and urea.

### Kinetic characterization of ammonia and urea oxidation

To determine whether N preference among these AOM species was associated with different cellular kinetics, we characterized ammonia- and urea-dependent oxidation kinetics using microrespirometry (Supplementary Table 2, Supplementary Discussion). Ammonia and urea-dependent oxygen consumption followed Michaelis-Menten kinetics in all AOM investigated here (Extended Data Fig. 11). Whereas the marine AOA *N. piranensis* showed similar apparent half-saturation constants (*K*_m(app)_) for ammonia and urea (0.70 ± 0.22 and 1.65± 0.82 µM, respectively), the soil AOA *N. viennensis* had comparable *K*_m(app)_ for ammonia (0.68 ± 0.16 µM), but significantly higher *K*_m(app)_ for urea (8.97 ± 1.27 µM) (Fig. 4a). Correspondingly, the specific affinities (*a*°) for ammonia and urea in *N. piranensis* and ammonia in *N. viennensis* were similar, whereas *N. viennensis a*° for urea was approximately 10-fold lower (Fig. 4b). The comammox *N. inopinata* showed comparably high affinity for ammonia as AOA species. However, unlike the AOA species, *N. inopinata* also showed a very high *a*° for urea, which exceeds that of *N. piranensis* and *N. viennensis* by more than 20- and even 90-fold, respectively (Fig. 4b, Supplementary Table 2). *N. inopinata* contains an ABC-type urea transporter (Extended Data Fig. 2) that may support its high affinity for urea via ATP-dependent urea uptake. Notably, the β-AOB *N. lacus* and *N. multiformis* had 2.7–6.2-fold higher *a*° for urea than for ammonia (Fig. 4b, Supplementary Table 2), comparable to or even higher (*P* < 0.05) than that of *N. viennensis* (Fig. 4b), suggesting some AOB common in soils and freshwater sediments might effectively compete with AOA for urea in these environments.

**Figure 4.**
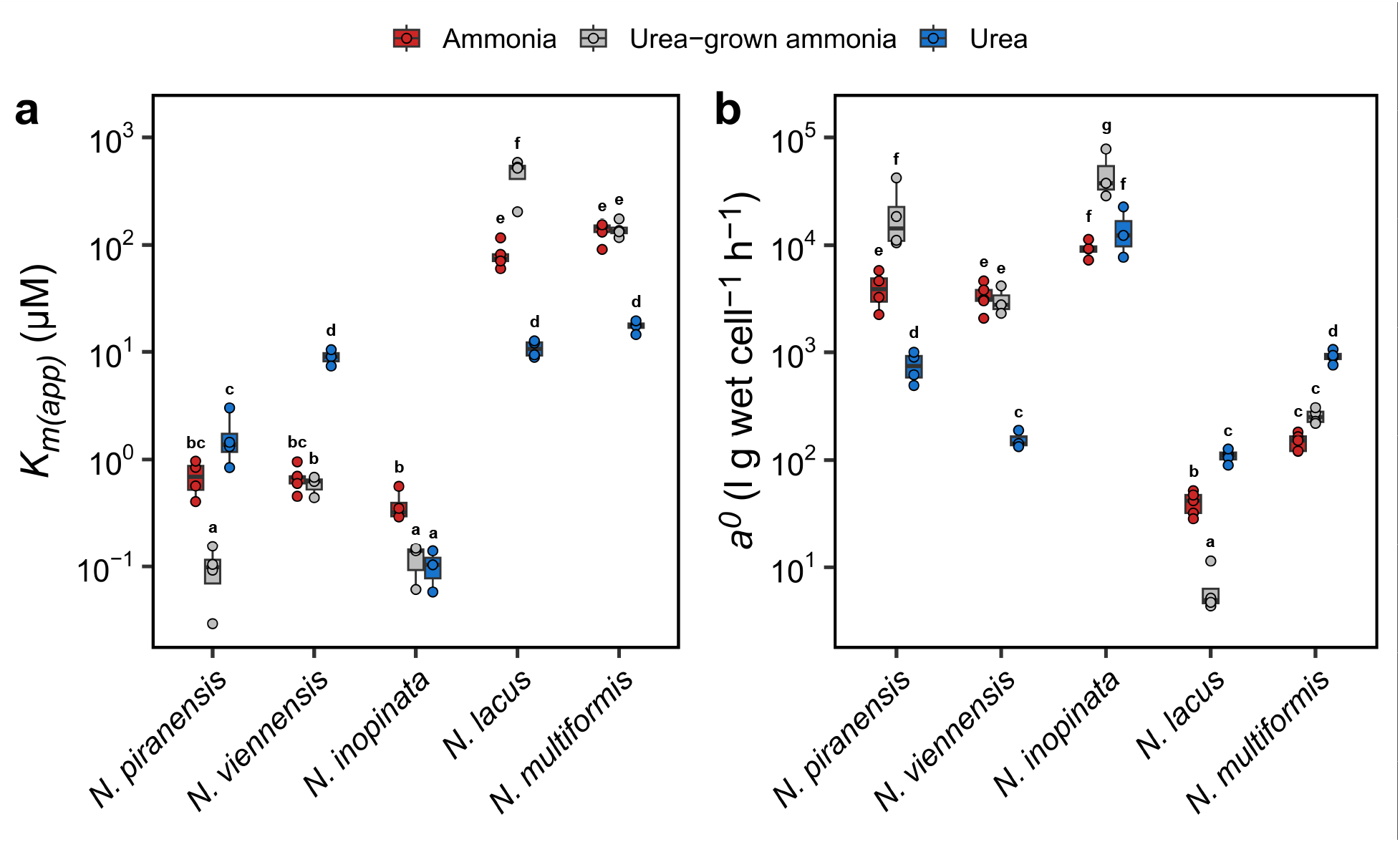
Michaelis-Menten kinetic constants of ammonia- and urea-dependent oxidation. (**a**) Half saturation constant (*K*_m(app)_), (**b**) specific substrate affinity (*a*°). *K*_m(app)_ and *a*° for ammonia were determined both in ammonia- (red) and urea-grown cultures (grey), and *K*_m(app)_ and *a*° for urea were determined in urea-grown species (blue). The upper limit, middle line, and lower limit of the boxplots indicate the 25th, 50th (median), and 75th percentile, respectively. Dots represent each replicate. Different lower-case letters denote statistically different groups based on Tukey’s HSD (*P* < 0.05).

We further tested ammonia affinities of urea-grown strains. To our surprise, only *N. viennensis* showed similar *K*_m(app)_ and *a*° independent of prior growth on ammonia or urea, whereas affinity for ammonia was significantly higher (*P* < 0.05) in urea-grown cells of *N. piranensis* and *N. inopinata* than ammonia-grown cells (Fig. 4), demonstrating a capacity by marine AOA and comammox to adapt kinetic properties. Such unexpected increase in affinity may prepare them for efficient uptake of their primary N source once it is again available. Taken together, the kinetic characteristics of AOM species track their N substrate preferences. *N. inopinata* is a notable exception, likely achieving unusually high urea affinity through ATP-dependent transport, which might allow comammox to successfully compete for both the preferred and alternative N sources in selected environments.

### Transcriptional patterns correlated with growth on single and mixed N substrates

Growth kinetics suggested that the differential use of ammonia and urea was coordinated by specific regulatory systems. Significant changes in gene transcription were correlated with the transition of the marine AOA *N. piranensis* and soil AOA *N. viennensis* to growth on their alternative N substrates (Fig. 5a, b). Cells were characterized at mid-exponential phase on a single N source (T1) and at 24 hours after introduction of the alternative N substrate (T2) (Fig. 5a, b). Additional samples were taken during the extended lag in *N. piranensis* growth during its transition from ammonia to urea (AU-T3) and after its consumption of the newly supplemented N species (AU-T4 & UA-T3) (Fig. 5a).

**Figure 5.**
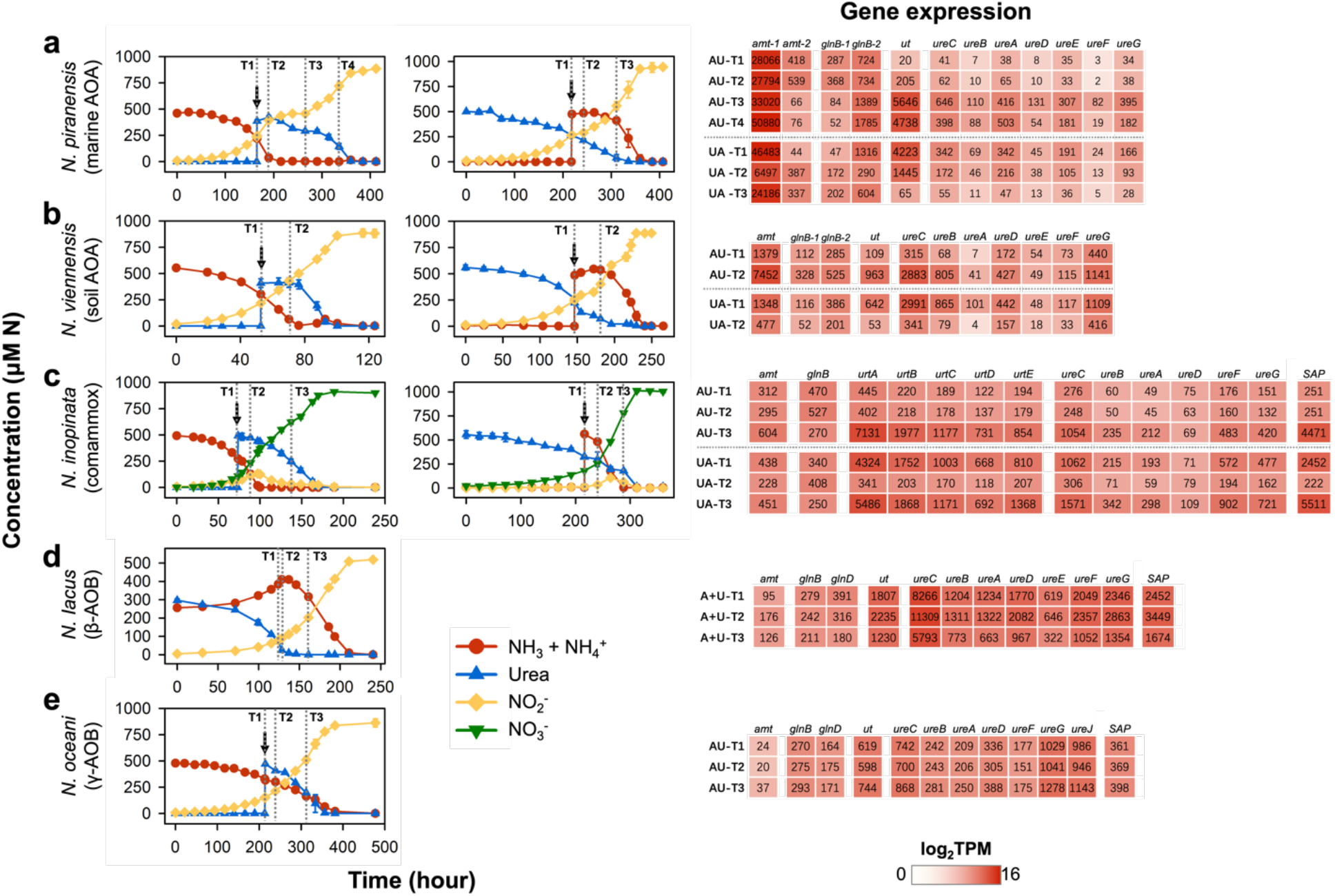
Transcriptional response of AOM species to substrate switch from singe N source to a mixture of ammonia and urea. The transcriptional changes of ammonia and urea-utilizing genes were investigated 24 h after addition of an alternative N substrate to exponential phase ammonia-grown or urea-grown (**a**) *N. piranensis*, (**b**) *N. viennensis*, (**c**) *N. inopinata*, (**d**) *N. lacus*, and (**e**) *N. oceani*. The growth data showing the times at which the alternative N substrate was added (arrows) and when cells were collected for transcriptomics analysis (dashed lines, T1−T4) are displayed in the left panels. T1 and T2 represent the sampling time points before the alternative N substrate addition and at ∼24 hours after addition, respectively. AU = ammonia spiked urea; UA = urea spiked ammonia; A+U = ammonia+urea. Markers and error bars on growth curves represent the average and one standard deviation of biological triplicates. The transcription data for key ammonia and urea-utilizing genes are profiled in the right panels. The average transcripts per million (TPM) values for each gene are shown in colored boxes and the color scale bar represents the log_2_(TPM). Gene full name and abbreviation pairs and source data are provided in Supplementary Tables 3 and 4.

In characterized AOA species, ammonia is used as a source of both energy and electrons through its oxidation by AMO, and for biosynthesis through the activities of glutamate dehydrogenase (GDH) and glutamine synthetase (GS) ^15, 25^. Transcription of *amoA* (AMO alpha subunit), *gdh*, and *gs* was relatively unchanged during exponential growth of *N. piranensis* and *N. viennensis* on either N substrate (AU-T1 vs. UA-T1, Supplementary Table 3). In contrast, the transcription of *amt1* (putative high-affinity ammonium transporter)*, ut* (urea transporter)*, ure* (urease), and *glnB* (P_II_ protein) were highly responsive to the addition of the alternative N sources (Fig. 5a, b).

Initially low transcription of the *ure* and DUR3-type *ut* (NPIRD3C_1395) in ammonia-grown *N. piranensis* increased to very high levels after switching to urea consumption (Fig. 5a). In turn, *ut* transcription was significantly depressed upon addition of ammonia to cultures growing on urea (UA-T2 vs. T1, *P* = 1.01 × 10^-13^), dropping to its lowest levels when urea was close to depletion (UA-T3). The *amt1* of *N. piranensis* was constitutively transcribed at high levels across all growth phases and conditions (Fig. 5a). Notably, it was most highly transcribed when growing on urea (46,483 TPM, Transcripts Per Million, UA-T1) and after transitioning to exponential growth on urea (50,880 TPM, AU-T4). Transcript abundance dropped more than 7-fold within 24 hours after addition of ammonia to urea-grown cultures (6,498 TPM, UA-T2). The increased transcription of *amt1* in urea-grown *N. piranensis* (46,483 TPM, UA-T1) is consistent with its higher specific affinity for ammonia, relative to ammonia-grown cells (28,066 TPM, AU-T1), which were determined by our kinetics analysis (Figs. 4).

Unlike *N. piranensis*, no differential transcription (*P* = 0.84) of *amt1* (NVIE_002420) was observed between the exponentially growing *N. viennensis* on ammonia (1,379 TPM, AU-T1) and urea (1,348 TPM, UA-T1), likely reflecting its comparable affinity for ammonia measured using either ammonia- or urea-grown cells (Fig. 4). The transcripts of a *ut* (NVIE_014780) and *ureC* (urease alpha subunit, NVIE_014740) located within the same operon (Extended Data Fig. 2) were ∼10-fold higher or lower within 24 hours following addition of urea or ammonia, respectively (AU- & UA-T2 vs. T1).

Distinct from the AOA species, the comammox *N. inopinata* exhibited much less stringent control of urea consumption. Urea was immediately hydrolyzed upon its addition to ammonia- growing cells (Fig. 5c). As observed for the AOA species, *N. inopinata* significantly increased (*P* < 0.01) the transcription of its single *amt* when transitioning to exponential growth on urea upon ammonia depletion relative to growing on ammonia (AU-T3 vs. T1), while its transcript abundance was significantly lower (*P* = 6.5 × 10^-27^) at 24 hours after ammonia addition to urea-grown cells (UA-T2 vs. T1). Its significantly higher transcription (*P* = 2.8 × 10^-3^) in urea-grown (438 TPM, UA-T1), versus ammonia-grown (312 TPM, AU-T1) cells is consistent with the increased ammonia affinity observed for cells precultured on urea (Fig. 4). Although the *urt* (urea transporter) and *ure* (NITINOP_3355–NITINOP_3362) were transcribed at significantly higher levels (*P* = 7.0 × 10^-25^) during exponential growth on urea (UA-T3 vs. T1), the cells maintained relatively high transcription levels of these genes when grown on ammonia (Fig. 5c), which is distinct from their extremely low transcription in the marine AOA *N. piranensis* before ammonia depletion (Fig. 5a). A gene encoding the putative outer-membrane short-chain amides/urea porin (SAP, NITINOP_3364) was found adjacent to the *ure/ut* operon and exhibited similar transcription patterns as *ut* (Fig. 5c), suggesting the regulation of urea uptake occurs in the transport systems of both the outer and inner membranes in comammox.

Since the substantial free ammonia released during consumption of urea by *N. lacus* would confound the effect of the added ammonia to urea-grown cultures, we instead examined transcript abundance at three time points of growth on an ammonia-urea mixture – exponential growth on urea (A+U-T1), near urea depletion (A+U-T2), and after shifting to exponential growth on ammonia (A+U-T3) (Fig. 5d). All genes involved in urea uptake and hydrolysis, including a putative *sap* (EBAPG3_007760), encoded in a single operon (EBAPG3_007760– EBAPG3_007800), were constitutively transcribed at relatively high levels at all time points, even when cells were using ammonia (Fig. 5d). This indicated that even while growing on ammonia, β-AOB *N. lacus* allocates substantial resources to facilitate the use of its preferred N source urea.

Unlike the highly variable transcription of *amt* observed in the AOA and comammox, a single *N. lacus* Rhesus-type *amt* was transcribed at constantly low levels in *N. lacus* (Fig. 5d), as has also been observed in *Nitrosomonas* β-AOB and *Nitrosococcus* γ-AOB species ^26, 27^. Of two *glnB*, one (EBAPG3_001140) was transcribed during all growth phases, whereas a paralog (EBAPG3_010980) showed very limited transcriptional activity. GlnB-type P_II_ proteins (such as GlnK in γ-proteobacteria) can control the uptake of extracellular ammonia through interaction with AMT ^28^. In well characterized heterotrophs, the transport of ammonia by the AMT can be blocked by binding to the GlnB protein. Binding to GlnB is controlled by its post-translational modification by a second regulatory protein (GlnD) via uridylylation/deuridylylation reactions in response to variations in intracellular energy and N status ^28–30^. A *glnD* homolog in *N. lacus* (EBAPG3_003255) suggests this organism encodes a similar regulatory system for N uptake. *GlnD* was transcribed at significantly higher levels (*P* = 6.6 × 10^-15^) when ammonia utilization was severely repressed (A+U-T1 and T2), suggesting the deuridylylated GlnB binds to AMT and blocks the transport of ammonia in the presence of urea as was demonstrated by our isotope tracking experiments (Fig. 2).

Given the extremely slow growth of the marine γ-AOB *N. oceani* on a urea-only medium (Fig. 1g), our transcript analysis only followed the response of cells growing on ammonia with subsequent urea addition. Urea hydrolysis started immediately upon addition and had no substantial effect on the growth rates or the transcription of genes directly involved in N species consumption (Fig. 5e). The lack of major differential transcription in *N. oceani* most likely reflects constitutive systems for ammonia and urea uptake, co-utilized without preference or adaptive lag (Fig. 5e).

## Discussion

The AOM in our study set displayed a wide spectrum of adaptive strategies to coordinate growth on a mixture of urea and ammonia. The tested AOA and comammox species preferentially used ammonia in contrast to β-AOB species, which favored urea. The characterized γ-AOB species co-utilized ammonia and urea as confirmed by lack of repression of the utilization of either N source. The preferential use of ammonia is likely a common lifestyle shared among *Nitrosopumilus* and *Nitrososphaera* species, as the characterized genomes of these two AOA clades all encode the putative high-affinity AMT and GlnB-type P_II_ proteins (Extended Data Fig. 1), which are suggested to be associated with efficient ammonia uptake and the repression of urea utilization in AOA, respectively (Supplementary Discussion). Likewise, the genes encoding P_II_ proteins (GlnB) and associated regulatory proteins (GlnD), which appear to be involved in the repression of ammonia usage in β-AOB, are universally present in *Nitrosospira* genomes, suggesting the preference for urea over ammonia may be common among ureolytic *Nitrosospira* species. In contrast, the *glnB* and *glnD* genes are largely absent in most *Nitrosomonas* genomes, which may in part explain the observed weaker repression of ammonia utilization by *Nitrosomonas ureae* compared to *Nitrosospira* species (Fig. 1).

Distinct lineages of AOM are generally thought to compete for a single N resource (ammonia), and differences in ammonia affinity are suggested as the main determinant of their competition and niche differentiation ^6–8^. However, different groups of AOM are often found to co-occur in terrestrial and estuarine environments ^1, 31–34^. Our study suggests that the differences in N substrate prioritization may mitigate direct competition. For example, β-AOB with a high affinity for urea could prioritize their growth on urea (Fig. 4) while *Nitrosopumilus*/*Nitrososphaera* AOA would thrive in the ammonia-depleted environment due to their much higher affinity for ammonia.

The biogeochemical role of urea in coastal ecosystems heavily impacted by runoff from fertilized agricultural areas and other anthropogenic sources has been recognized ^21, 22^. In order to compete with many other microbial groups for urea ^35^, marine AOA would need to respond rapidly to periodic enrichment of urea in estuarine and coastal regions ^21^. However, we observed a long diauxic lag in transition from ammonia to urea utilization in *N. piranensis*, a coastal AOA species (Figs. 1a, 5a, Extended Data Fig. 12b, Supplementary Discussion). We can now postulate that competitive advantage among *Nitrosopumilus* and related AOA is derived not solely from use of both N substrates, but also from the adaptive capacity of marine AOA to increase their affinity for ammonia upon ammonia exhaustion (Fig. 4), resulting in more efficient scavenging of ammonia released from organic matter decomposition by other microorganisms^35, 36^.

One of the most significant findings of our study is that ammonia oxidation and ammonia assimilation are both transport-dependent. Our stable isotope tracking analysis clearly demonstrated that β-AOB did not oxidize extracellular ammonia when growing on a mixture of ammonia and urea, rather only ammonia originating in the cytoplasm from urea hydrolysis was oxidized. Ammonia transport is blocked when N needs are satisfied by cytoplasmic hydrolysis of urea, most likely by binding of a GlnB-type P_II_ regulatory protein to the AMT. Thus, these data also implicate an undescribed mechanism for direct or facilitated delivery of cytoplasmic ammonia to the AMO active site.

Since extracellular ammonia must be transported into the cytoplasm by the AMT before being oxidized by the AMO, the AMT must be the rate-limiting functional unit that determines affinity for both the assimilation and oxidation of ammonia among AOM. Recently experimentally determined *K*_m_ of ammonia transporters (e. g. ∼800 µM for *E.coli* amt, 3.9 mM at pH=7 for *Archaeoglobus fulgidus* Amt1 ^37–39^) are higher than the cellular *K*_m(app)_ determined in our study. However, kinetic data on ammonia assimilation by phytoplankton or heterotrophic bacteria widely believed to be conferred by Amt-type transporters indicates that ammonia transporter variants with higher affinities (*K*_m(app)_ 0.1–10 µM) exist in the environment ^40, 41^. In our study, a 5.1-fold increase in the *a*° of *N. piranensis* for ammonia was correlated with increased transcription of its high-affinity AMT (Figs. 4b and 5a). In contrast, the transcription of *amo* remained relatively unchanged during exponential growth of *N. piranensis* on either N substrate (AU-T1 vs. UA-T1, Supplementary Table 3). Combined modelling and substrate uptake studies of oligotrophic marine bacteria have demonstrated the relationship between the number of cell surface transporters and *a*° ^42, 43^. A recent extensive investigation of apparent affinity of different AOM lineages showed that the affinities for ammonia correlate with their cell surface area to volume ratios ^8^, suggesting that a higher cellular affinity for ammonia is achieved by the higher number of AMTs that can be accommodated on a larger surface area. Since it is unlikely that single species can substantially change its cell surface area to volume ratio, the increased cellular ammonia affinity observed for urea-grown *N. piranensis* and *N. inopinata* is most likely caused by the increase in the density of high-affinity AMT per cell surface area.

Unlike other AOA lineages, *Nitrosocosmicus* species lack the putative high-affinity AMT and exhibit highly active urea hydrolysis activity when grown in urea-only media ^44^, which suggests that they may employ a distinct regulation pattern for N substrate utilization compared to *Nitrosopumilus* and *Nitrososphaera* species. In addition, while both clade A and clade B comammox species possess ATP-dependent urea transporters, they differ in their encoded ammonia transporter types and the presence of GlnD proteins ^45^ (Extended Data Fig. 1). As more diverse comammox strains become available in the future, it will be interesting to see whether different transporter inventories impact the regulatory mechanisms controlling N catabolite repression in comammox.

In summary, our study revealed an unexpected diversity in substrate preference and supporting regulatory systems among AOM and has significantly changed our understanding of the ecophysiology and niche differentiation within this globally abundant functional guild. Our surprising finding that cellular ammonia affinity is not an intrinsic property of AOA and comammox species, but a property that can be adaptively regulated through transport in response to the environmental ammonia availability, has implications on how N resource competition is modelled among AOM groups. Adding these novelties to models will ultimately offer a more holistic and accurate representation of N resource partitioning through cooperative division of labor among AOM in natural and engineered systems.

## Acknowledgements

We thank Zach Perry, Bella Quan, and Trevor Tubbs for technical assistance in growth experiments. We appreciate the work of Sergey Oleynik in maintaining the mass spectrometer for stable isotope analysis. *N. inopinata* was generously provided by Dr. Holger Daims, University of Vienna. *N. piranensis* was generously provided by Dr. Alyson Santoro and Dr. Barbara Bayer, University of California, Santa Barbara. *N. multiformis* was kindly provided by Dr. Jeanette Norton, Utah State University. *N. ureae* was kindly provided by Dr. Andreas Pommerening-Röser, University of Hamburg. This work was supported by the National Natural Science Foundation of China grants 42277304 and 41977056 (to BZW), 42276117 (YZheng), and 42125603 (to YZhang). This work is also supported by the startup funding of the University of Oklahoma (to WQ). Work was also funded by the US Department of Energy’s Office of Science, Division of Biological and Environmental Research grant Program (Grant DE- SC0020356) (to MKHW, XM, DAS, CP, ZF, BA, SPW) and by the Defense Advanced Research Projects Agency (Contract Number: HR0011-17-2-0064) (to MKHW, DAS, SPW). Funding was also provided by the Florida Agricultural Experiment Station Hatch project FLA-FTL- 005680, UF IFAS Early Career Award, and USDA NIFA award #2022-67019-36501 (to WMH). Part of this work was performed at Lawrence Livermore National Laboratory under Contract DE-AC52-07NA27344. The participation of XW and BBW was supported by Simons Foundation grant 675459 to BBW. XS was supported by G. Evelyn Hutchinson postdoctoral fellowship from Yale Institute for Biospheric Studies at Yale University. HU was supported by the National Science Foundation grant (DEB-1664052).

## Author contributions

WQ, DAS, BBW, XM, WMH, and MKHW designed the experiments. WQ, SPW, YZ, EC, XL, JJ, XW, BA, and ZF performed the experiments and analyzed data with input from BW, HL, LH, QT, WWC, XS, MW, LN, and KAH. WQ, SPW, DAS, WMH, and MKHW wrote the manuscript, with contributions and approval from all other authors.

## Data and materials availability

All data are available in the main text or the supplementary materials.

Supplementary Information is available for this paper.

Correspondence and requests for materials should be addressed to Wei Qin, Willm Martens-Habbena, and Mari Winkler.

Reprints and permissions information is available at www.nature.com/reprints.

